# Coupling between Notch signalling and junctional mechanics during asymmetric division of sensory organ precursors

**DOI:** 10.64898/2026.07.10.737684

**Authors:** Mathieu Pinot, Roland Le Borgne

## Abstract

Mechanical forces and signaling pathways are increasingly recognized as interdependent regulators of epithelial morphogenesis, yet their combined role in cell fate acquisition remains poorly understood. Here, we investigate the interplay between adherens junction mechanics and Notch receptor signaling during the asymmetric division of sensory organ precursors in the *Drosophila* pupal notum epithelium. Using quantitative live imaging and laser ablation, we identify the newly formed interface between SOP daughter cells as a mechanically specialized junction, characterized by persistently low membrane tension, distinct adhesive organization, and a unique cortical actomyosin architecture.

We propose that low membrane tension may facilitate efficient Notch activation, as ligand-mediated endocytosis promotes Notch signaling by generating traction forces of a few piconewtons, oriented perpendicular to the plasma membrane. Perturbations of Notch pathway activity systematically alter junctional recoil following laser ablation, with reduced Notch signaling correlating with increased tension. Conversely, constitutive Notch activation in a *Notch* loss-of-function context is sufficient to restore a low-tension state. These findings suggest that Notch signaling actively shapes the mechanical properties of its signaling interface, indicating reciprocal interactions between mechanics and signaling.

Together, our results support a model in which Notch activity and junctional mechanics are coupled during asymmetric cell division, highlighting how local mechanical states may contribute to the robustness of cell fate specification in epithelia.

## Introduction

An emerging body of evidence suggests that epithelial tissues integrate mechanical forces with signalling pathways to coordinate cell fate decisions, tissue patterning, and morphogenesis (1, 2). Mechanical forces are now recognized as critical regulators of Notch signalling, acting through two distinct but complementary mechanisms. Firstly, ligand-mediated forces arise from the endocytosis of Notch ligands, generating a traction force oriented perpendicular to the plasma membrane. This force promotes the conformational changes necessary for proteolytic cleavage of Notch, enabling the release of its intracellular domain for nuclear translocation and activation of target genes (3–5). Secondly, the mechanical forces at the tissue level, exerted by the surrounding epithelial cells in the form of tensile and compressive forces at the level of the adherens junctions (AJs), as well as shear stresses, shape the cellular geometry and influence the distribution and activity of signalling receptors at cell-cell interfaces (6). Actomyosin contractility, adhesion dynamics, and membrane tension contribute to this mechanical regulation. However, how these inputs collectively influence the acquisition of cell fate remains largely underexplored. Notch signalling, which relies on direct cell–cell contact via *trans*-interaction of its receptor with ligands, is involved in intra-lineage signalling to regulate binary cell fate acquisition and lateral inhibition to control patterning. The interplay between mechanical forces and Notch signalling is a critical regulator of lateral inhibition, operating at both global and local scales. Global forces can spatially and temporally restrict Notch activation to specific tissue domains, as in the mammalian inner ear, while local epithelial cell-mediated forces drive symmetry breaking and collaborate with Notch to refine patterns in intestinal organoids (6) and the zebrafish myocardial wall (7). Moreover, Notch-mediated lateral inhibition feeds back to modulate intra-and inter-cellular mechanics, generating complex patterns like those in the *Drosophila* retina (8). Despite these observations, the molecular mechanisms underlying this coupling remain elusive. For example, how mechanosensitive pathways regulate Notch activity, and how Notch reciprocally controls cell adhesion, morphology, and intracellular forces are unresolved. Whilst the links between mechanical forces and the Notch signalling pathway are beginning to be elucidated, it is not yet known whether, and if so how, the mechanical forces generated by epithelial cells influence the Notch-dependent binary fate decision amongst the daughter cells of a progenitor cell.

The notum of *Drosophila melanogaster* is a proliferative monolayered epithelium composed of two cell populations, epidermal cells (ECs) which divide symmetrically and sensory organs precursors (SOPs). SOPs undergo asymmetric division to generate two daughter cells, pIIa and pIIb, which adopt distinct fates through differential activation of Notch (9, 10)The newly formed pIIa–pIIb interface constitutes a highly specialized signalling platform where Notch, its ligand Delta, and associated cell polarity and membrane trafficking regulators including Par3, Sanpodo and Neuralized assemble into clusters (11). Two pools of Notch, located apically and basally to the midbody contribute to Notch signalling during SOP cytokinesis, with the basal pool being the main contributor (12, 13). Although molecular mechanisms governing Notch activation at this interface have been characterized, the mechanical properties of this signalling interface and its influence on fate acquisition remain poorly understood. In particular in contrast to the basally located Notch signalling clusters, the apical clusters are present in the plane of AJ, potentially prefiguring distinct local mechanical constrains (11). Indeed, epithelial cells can modulate junctional tension through remodelling of E-cadherin organization, actomyosin contractility, and vesicular trafficking. However, whether mechanical properties of the pIIa–pIIb interface influence Notch activation, or conversely whether Notch signalling actively shapes the mechanical state of this interface, is unknown.

Here, we combine quantitative live imaging, laser ablation, and genetic perturbations to characterize the mechanical properties of SOPs and their daughter cells. We find that the pIIa–pIIb interface exhibits persistently low membrane tension and a distinct adhesive and cortical organization. We show that manipulating Notch pathway activity systematically alters junctional recoil following junction ablation, and forced activation of Notch is sufficient to impose a low-tension state. Our results demonstrate that Notch signalling actively instructs the mechanical properties of the daughter cell interface, revealing a coupling between signalling and mechanics during asymmetric cell division.

## Results

### SOPs exhibit distinct apical geometry and actomyosin organization

SOPs and ECs have clearly distinct characteristics in relation to the curvature of the bicellular junction. Namely, in the AJ plane marked with E-Cadherin::GFP (Ecad::GFP), EC cells have a regular polygonal shape, whilst SOP cells have a plasma membrane that curves inwards, giving them a characteristic star-shaped morphology (Fig. 1A) (14). Quantification of the shape morphology revealed a reduced area-to-perimeter ratio in SOPs compared to ECs at the AJ plane (Fig. 1A,A’; Fig. S1A’). This difference was not detected at the level of septate junctions (SJs) or in more basal planes (Fig. 1A’, Fig. S1A’’A’’’), indicating that the shape control is restricted apically. Because AJs are mechanically coupled to actomyosin contractility (15), we examined the distribution and dynamics of the regulatory light chain of nonmuscle type 2 myosin, hereafter referred to as MyoII. SOPs displayed a denser and a thicker medial MyoII network than ECs (Fig. 1B-B’’)(14, 16). Time-lapse imaging revealed a pulsatile MyoII behavior (Fig. 1C, C’) in both cell types with a similar pulse frequency (∼230 +/- 129 seconds for ECs ; ∼220 +/- 104 seconds for SOPs ; Fig. 1C’), but with a significantly greater pulse amplitude in SOPs (∼1.5-fold magnitude; Fig. 1C’’, lower panel). Additionally, medial MyoII flows toward the cortex preceded SOP membrane deformation (Fig. S1B), consistent with contractility-driven curvature and a SOP-specific mechanical coupling. To directly assess junctional tension, we performed two-photon laser ablation at the AJ plane. EC–EC junctions exhibited rapid recoil (30–45 seconds) followed by MyoII-dependent repair (Fig. 1D; Fig. S1CC’)(17). In contrast, SOP–EC junctions showed ∼6-fold lower recoil velocity and ∼7-fold lower final displacement (Fig. 1D–D’), indicating a reduced junctional tension. No relaxation was observed at the SJ plane, marked with ATPa::YFP (Fig. S1D,D’), confirming the localization of tensile forces in the AJ plane (15).

**Figure 1.**
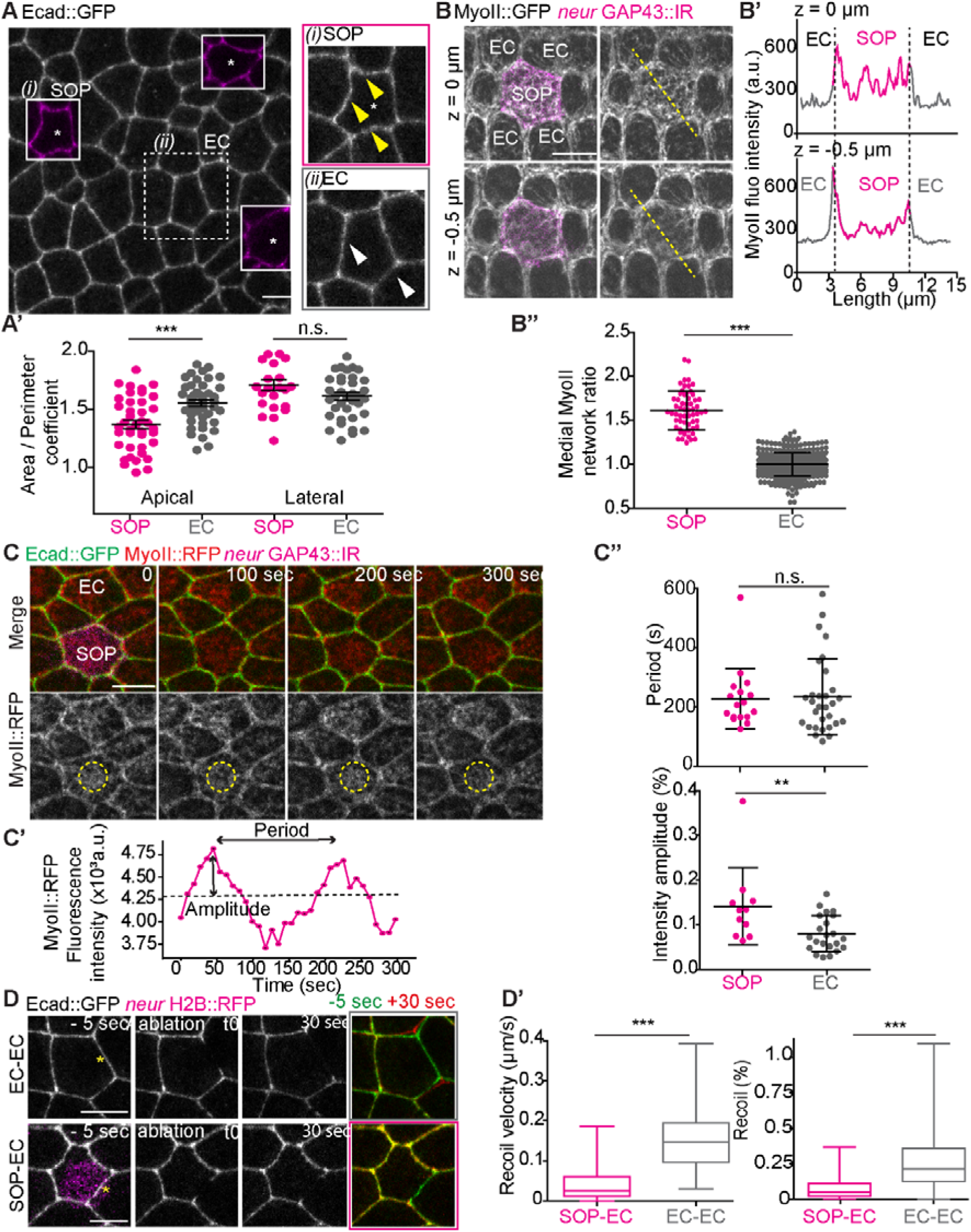
Interplay between atypical shape and mechanical properties of sensory organ precursors (SOPs). (A) Confocal image of the *Drosophila* pupal notum epithelium at the adherens junction (AJ) level, labelled with Ecad::GFP. SOPs are identified by the nuclear marker H2B::RFP expressed under the *neuralized* promoter (magenta). Dashed squares indicate (i) SOP and (ii) epidermal cells (ECs). Yellow arrows point to curved junctions in SOPs; white arrows indicate straight junctions in ECs. (A’) Distribution of the cell area to perimeter ratio measured at apical (n= 42 for SOPs and n= 71 for ECs) and lateral (n= 27 for SOPs and n =48 for ECs) levels for ECs (grey dots) and SOPs (magenta dots). (B) Localization of MyoII::GFP in SOPs at AJ level and 0.5 µm below. SOPs are labelled with the membrane marker GAP43 expressed under the *neuralized* promoter. (B’) MyoII intensity profile of MyoII::GFP along the yellow dashed line shown in (B). (B’’) Quantification of medial MyoII::GFP intensity in ECs (greys dots, n= 366) and SOPs (magenta dots, n= 59), normalized to the MyoII::GFP medial network intensity measured in three adjacent ECs (see Materials and Methods). (C-C’’) Oscillations of the medial MyoII network in SOPs. (C) Time-lapse imaging of SOP during interphase expressing Ecad::GFP (green) and MyoII::RFP (red and grey). (C’) Temporal profile of mean medial MyoII intensity marked with the ROI represented on C. (C’’) Quantification of oscillation period and amplitude of MyoII intensity in SOPs (n= 17) and ECs (n= 30). (D) Ablation of adherens junctions at EC-EC or SOP-EC interfaces. SOPs are marked with H2B::RFP expressed under *neuralized* promoter (magenta). Left and middle panels show cells before and 1 second after ablation, respectively. Right panels overlay images taken 5 seconds before ablation (green) and 30 seconds after ablation (red). (D’) Quantification of recoil velocity (left panel) and recoil final displacement (right panel) after ablation at EC-EC (n= 148) and SOP-EC (n= 30) interfaces. Recoil final displacement in % is defined by (_Lmax following ablation_ – L _before ablation_ / L _before ablation_) where L is the length of the junction. Yellow stars mark ablated junctions. Scale bars: 10 µm. Times in seconds.

Together, these data indicate that SOPs possess enhanced medial actomyosin dynamics with reduced junctional tension, revealing a distinct apical mechanical organization. These observations raise the question of whether these mechanical properties are transmitted to daughter cells pIIb and pIIa interface.

### The pIIa–pIIb daughter cell interface maintains low membrane tension

SOPs divide asymmetrically to generate a posterior pIIa cell and a smaller anterior pIIb cell (Fig. 2A,(10)). The formation of pIIa-pIIb bicellular junctions results in vertices that are very close to one another, with the distance between them being less than 0.5 µm (Fig. 2BB’; Fig S2A’), and remaining juxtaposed for at least 30 minutes after cytokinesis (Fig. S2A). Bicellular junctions of ECs and SOP daughters repair efficiently following laser ablation (Fig. S1CC’). However, recoil analysis revealed marked differences in tension between the two cases. EC junctions relaxed with a mean recoil velocity of 0.15 +/- 0.07 μm/s (Fig. 2C’, left panel), inversely correlated with initial junction length (Fig. 2C’). At comparable junction length (between 3 and 5 µm), EC recoil velocity (∼0.15 +/- 0.07 μm/s) and final recoil (25% of the initial length, Fig. 2C’ right panel) were ∼4-fold and ∼5-fold higher, respectively, than that of the pIIa–pIIb interface (V recoil∼0.05 μm/s and final recoil is ∼5%) (Fig. 2CC’).

**Figure 2.**
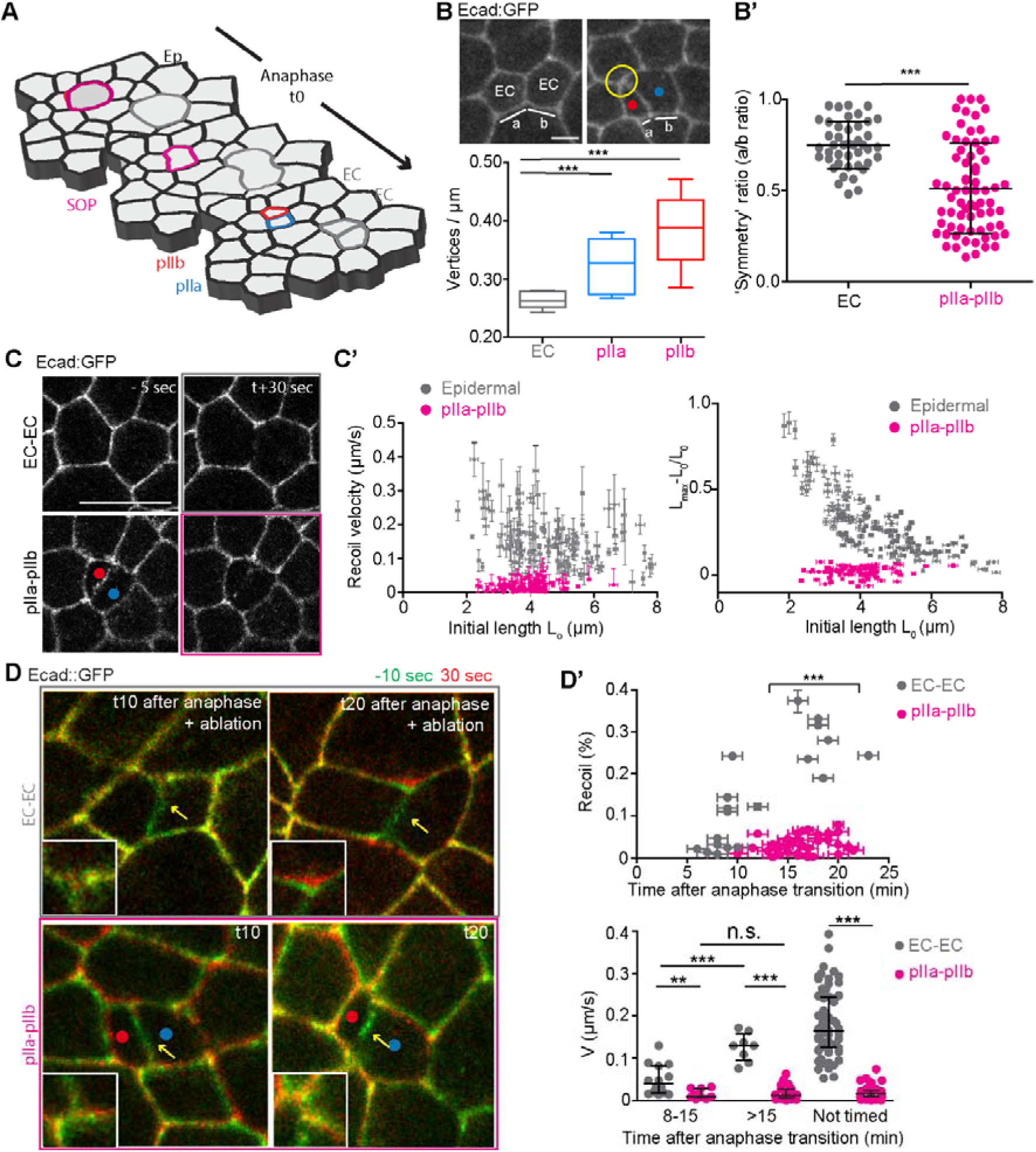
pIIa-pIIb adhesive interface exhibits low membrane tension. (A) 3D-schematic representation of the *Drosophila* pupal notum divisions composed of ECs (grey cells) and SOPs (magenta cells). t0 is considered as the metaphase to anaphase transition. SOP division give rise to pIIb (red dot) and pIIa cells (blue dot). (B) Geometrical distribution of vertices of the new formed EC-EC and pIIa-pIIb interface at 30 minutes after anaphase onset. (B) Quantification of vertices / µm in the case of EC (n= 26), pIIa and pIIb (n= 38) cell. (B’) Quantification of the symmetry ratio in EC (n= 52) and pIIa-pIIb (n= 76) junction. The symmetry ratio is defined as the ratio between the shortest length (a) and the longest length (b) between the new formed vertex and the closest vertex. A symmetry ratio around 1 means that the new junction is formed symmetrically. A symmetry ratio closed 0 means that two vertices are less than 0.5 µm apart (yellow circle, panel (B)). (C) Two examples of ablation of new EC-EC junction and new pIIa-pIIb interface. Left panels represent cells before ablation. Right panels represent cells 30 seconds after ablation. (C’) Quantification of the recoil velocity (µm/s) and the recoil maximal displacement (%) in function of the length before ablation Lo (n= 116 for EC-EC and n= 57 for pIIa – pIIb). Black dots represent EC-EC interface. Magenta dots represent pIIa-pIIb interface. (D) Ablation of EC-EC and pIIa-/pIIb interfaces performed 10 minutes and 20 minutes after anaphase onset. The images represent the merge of the cell taken 10 seconds before ablation (green) and 30 seconds after ablation (red). White squares represent magnification of a vertex. The separation of green and red channels means that the junction followed a relaxation process after ablation. Red and blue dots correspond to the anterior pIIb cell and the posterior pIIa cell, respectively. Yellow stars represent the ablated junction. pIIa/pIIb cells were identified with nuclear marker H2B::RFP expressed under the *neuralized* promoter. (D’) Quantification of recoil maximal displacement and recoil velocity following ablation between 5 and 25 minutes after anaphase onset (n= 21 for EC-EC and n= 39 for pIIa – pIIb). Black dots represent EC and magenta dots represent pIIa/pIIb junction. Scale bars are 5 µm. Times are expressed in minutes in (D) and in seconds in (C).

Time-resolved analysis (Fig.S2B) relative to anaphase onset showed that EC junctional tension increased between around 8 and 15 minutes after division (Fig. 2D’D’’). In contrast, recoil velocity at the pIIa–pIIb interface remained persistently low throughout this period (Fig. 2DD’). Similarly, pIIa–EC and pIIb–EC contacts displayed reduced recoil compared to EC–EC junctions (Fig. S2C). Thus, low membrane tension is a stable and specific property of SOP daughter interfaces in contrast to EC membrane interfaces.

### The pIIa–pIIb interface exhibits distinct adhesive and cortical organization

In order to determine how SOP daughter interface differs from a typical EC-EC interface and in addition to revealing that pIIa-pIIb interface exhibits a low membrane tension, we next examined the specific molecular architecture of the SOP daughter interface. Indeed, whereas in EC junctions, the laser ablation caused loss of Ecad signal along the entire bicellular junction, the loss of Ecad signal remained confined to the ablated region at the pIIa–pIIb interface (Fig. S3 A,A’), indicating differential adhesive behaviour. In addition, whereas EC junctions displayed homogeneous, apically restricted Ecad staining (∼0.5 μm thickness), the pIIa–pIIb interface showed discontinuous labelling extending ∼1 μm along the apico-basal axis (Fig. 3A,A’, Fig. S3 B,B’). FRAP analysis revealed slower recovery and a reduced mobile fraction of Ecad at the SOP daughter interface (Fig. 3B,B’). Consistent with reduced mechanical load, the Vinculin tension sensor was weakly and discontinuously recruited compared to EC-EC interface (Fig. S3C).

**Figure 3:**
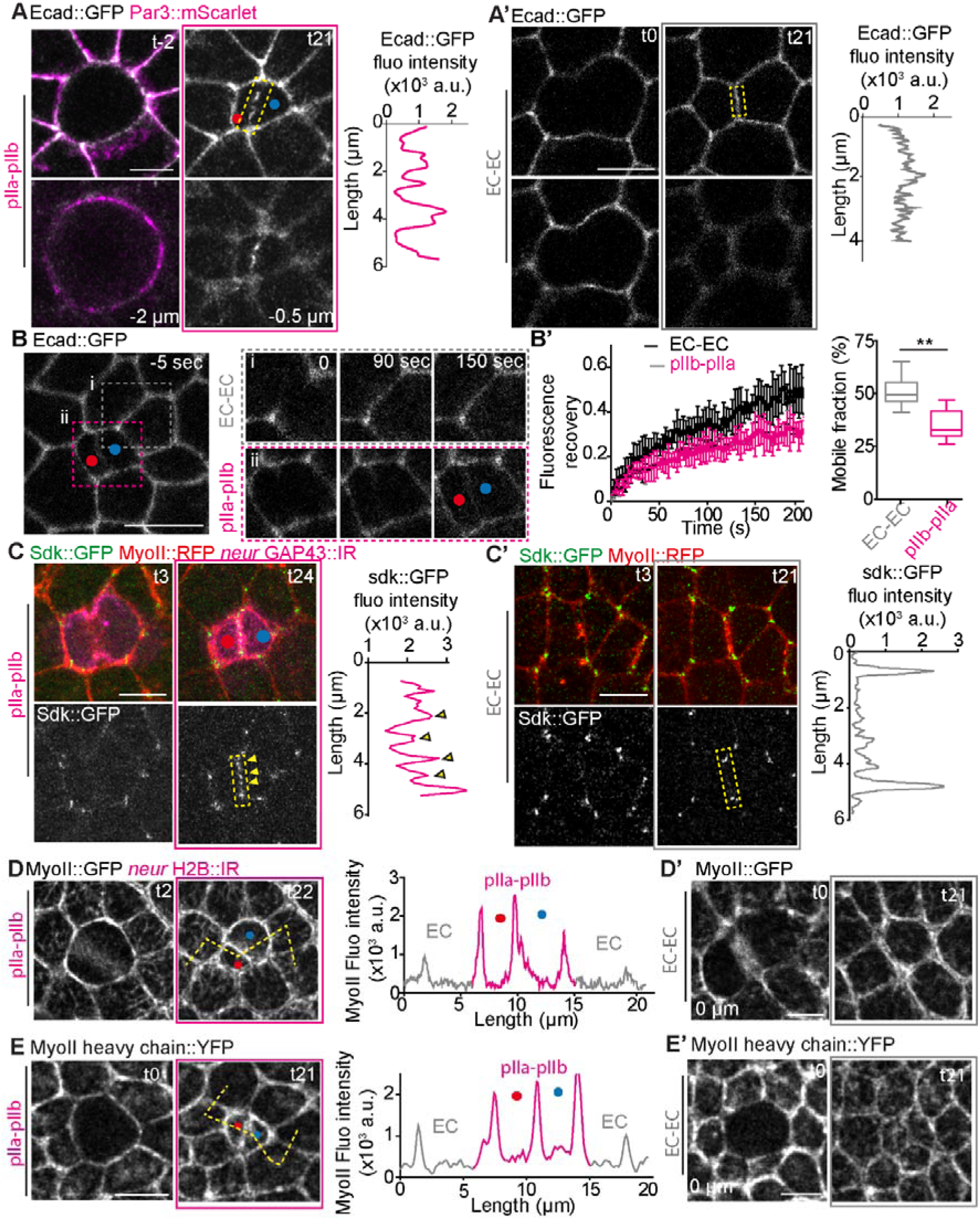
Distribution and dynamics of adhesive markers during SOP division. (A-B) Ecad::GFP imaging at times t0 and t21 during SOP cytokinesis and EC cytokinesis at AJ level and 0.5 µm below. (A’-B’) Ecad intensity plot profile of new formed interfaces. SOP daughter cells identified by exploiting the fact that Par3::mScarlet localizes asymmetrically. Yellow arrowheads mark Ecad::GFP discontinuities. (B-B’) FRAP experiments on Ecad::GFP at pIIa/pIIb and neighbouring EC interfaces. (B) Field of view of two photobleached junctions: (i) epidermal interface (upper), (ii) pIIb-pIIa interface (lower). Yellow dashed squares mark FRAP regions. (B’) Photobleaching events at EC and pIIa-pIIb junctions. (B-B’) Fluorescence recovery curves of Ecad::GFP at EC-EC (n = 8) and pIIa-pIIb interfaces (n= 9). (B’) Quantification of mobile fraction: 50 ± 15% at EC interfaces, 30 ± 15% at pIIa-pIIb interfaces. (C-C’) Imaging at different time points of Sdk::GFP and MyoII::RFP during SOP (n= 14) and EC (n= 15) cytokinesis. Intensity profiles at 21 minutes after anaphase onset along yellow dashed lines. Yellow arrowheads mark Sdk::GFP at the pIIa-pIIb bicellular junction; blue arrowheads mark Sdk::GFP at tricellular junctions. (D-D’) Imaging at different time points of MyoII::GFP during SOP and EC cytokinesis at AJ level. (G’-H’) Intensity profiles at 21 minutes after anaphase onset along yellow dashed lines, showing shell behaviour of pIIb-pIIa cells. (E-E’) Imaging at different time points of MyoII heavy chain::YFP during SOP (n= 10) and EC cytokinesis (n= 8) at AJ level. Intensity profiles at 21 minutes after anaphase onset along yellow dashed lines, showing shell behaviour of pIIb-pIIa cells. Scale bars: 5 µm. Times in minutes; seconds in (B’). *t*_₀_ corresponds to anaphase onset.

Sidekick (Sdk), a component of the tricellular junction that is normally enriched at tricellular junctions, is instead distributed in a punctate pattern along the entire length of the pIIa–pIIb interface (Fig. 3C, C’). In parallel, MyoII light chain and MyoII heavy chain revealed a thicker cortical MyoII network forming a shell-like structure in SOP daughter cells (Fig. 3DD’; Fig. 3EE’; Fig. S3DD’-EE’). Overall, these characteristics highlight the fact that the pIIa-pIIb interface differs from an EC-EC interface both mechanically and molecularly, with distinct organisation of adherens junctions and cortical contractility. Because this interface is the site where the Notch and Par3 clusters assemble and where the activation of the Notch signalling pathway occurs (11), we next examined whether its mechanical state is functionally linked to activation of the Notch pathway.

### Notch-regulatory trafficking modulates daughter cell interface tension

As the distribution of E-cadherin is discontinuous at the pIIa-pIIb interface (Fig. 2A), we sought to determine whether regulators of both Ecad trafficking and Notch signalling influence the mechanical properties of pIIa-pIIb interface. We chose the period between 15 and 30 min after anaphase onset due to specific mechanical properties described above (Fig; 2DD’) and Notch transient apical localisation (Fig. 4 A,A’). We first investigated the effect of Sec15 loss of function, a component of exocyst complex required for recycling of Ecad to plasma membrane and for the activation of Notch (18, 19). Loss of Sec15 did not alter recoil following ablation at EC junctions but induced a ∼4-fold increase in final displacement at the SOP daughter interface (Fig. 4 B,B’ and D,D’). Thus, in this experimental design, loss of Notch signalling is associated with elevated tension specifically in SOP daughter cells. Conversely, loss of AP-47 which regulates post-Golgi trafficking of E-Cad and negatively regulates Notch signalling (20–22), gave rise to a reduced recoil following ablation in both EC and SOP junctions (∼3-fold decrease in ECs and ∼2-fold decrease in SOP daughters; Fig. 4C, C’, DD’, Fig. S4 A,A’’). This is consistent with a decreased membrane tension, also associated with a characteristic tortuous interface in ECs and SOPs daughter cells junctions (21) (Fig. 4D; Fig. S4 A,A’).

**Figure 4:**
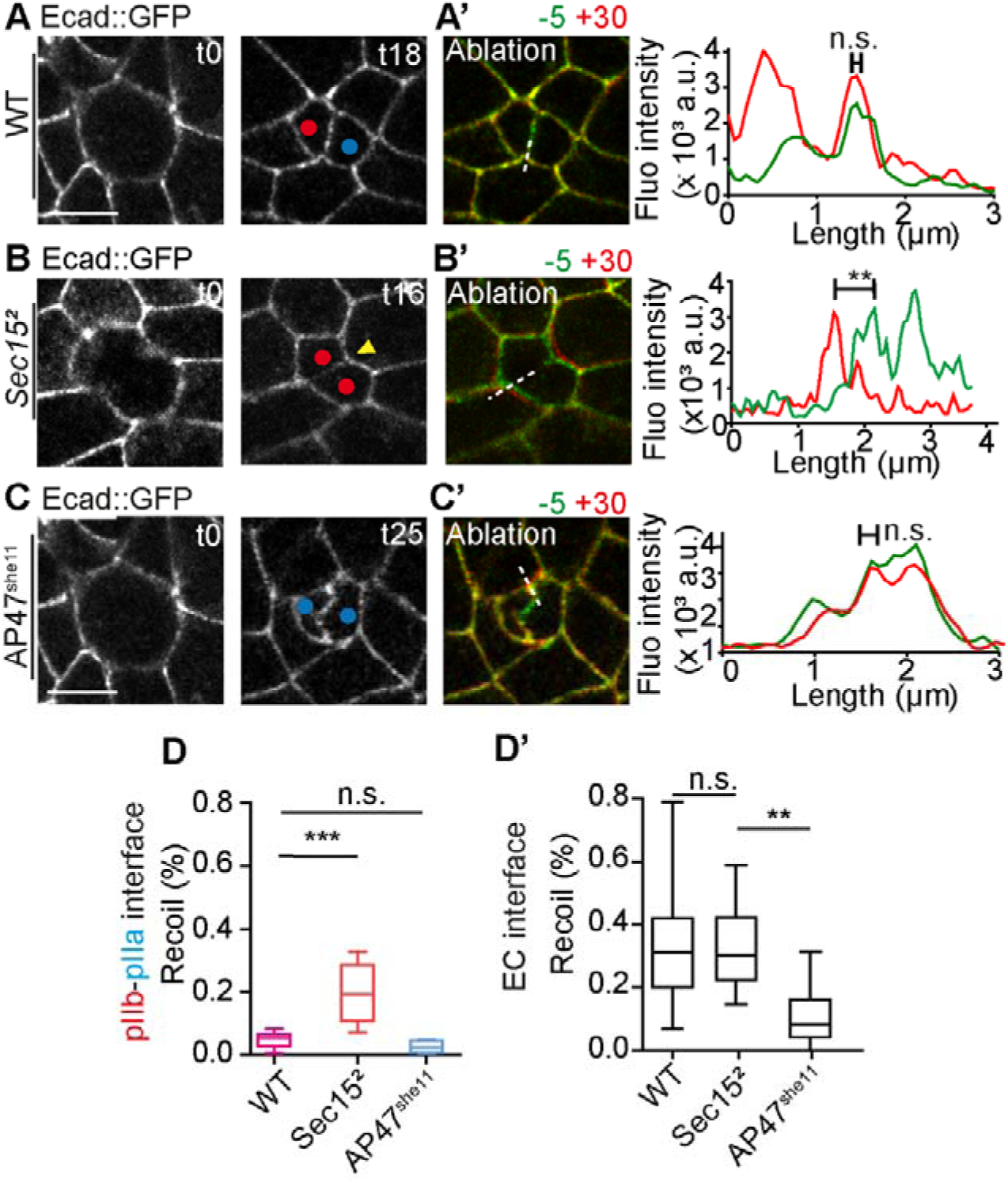
Role of vesicular trafficking regulators on the mechanical properties of the newly formed pIIa-pIIb interface. (A) Ablation of pIIa-pIIb interface at 18-minutes after anaphase onset. Left panel represents cell before ablation. Merge images show merges 5 seconds before ablation (green) and 30 seconds after (red). Fluorescence intensity profiles represent Ecad::GFP signal along the dashed white line before (green) and after ablation (red). (B) Localization of Ecad::GFP at t16 min after the anaphase onset upon Sec15 silencing. (C’) Ablation of pIIb-pIIb-like interface at 16 minutes. Images show merges 5 seconds before ablation (green) and 30 seconds after (red). Fluorescence intensity profile represents Ecad::GFP signal along the dashed white line before (green) and after ablation (red). (C) Distribution of Ecad::GFP at t25 min after the anaphase onset upon AP47 silencing. (right panel). (C’) Ablation of pIIa-pIIa-like interface at 25 minutes. Images show merges 5 seconds before ablation (green) and 30 seconds after (red). Fluorescence intensity profiles represent Ecad::GFP signal along the dashed white line before (green) and after ablation (red). (D-D’) Quantification of normalized recoil final displacement (L_₃₀_–L_₀_)/L_₀_ following ablation at 18 to 25 minutes after anaphase onset for (D) pIIa-pIIb (n= 16 for WT; n= 9 for Sec15^2^ mutant; n= 5 for AP47^she11^ mutant) and (n= 148 for WT; n= 24 for Sec15^2^ mutant; n= 33 for AP47^she11^ mutant) and (D’) EC-EC interfaces (n= 148 for WT; n= 24 for Sec15^2^ mutant; n= 33 for AP47^she11^ mutant). Red and blue dots represent anterior pIIb (or pIIb-like) and posterior pIIa (or pIIa-like) cells, respectively. pIIa/pIIb cells are identified with nuclear H2B::RFP under *neuralized* promoter. Scale bars: 5 µm. Times in minutes. *t*_₀_ = anaphase onset.

Thus, altogether these results show that high and low membrane tension of the SOP daughter cell interface are correlated with the loss and gain of the Notch signalling pathway, respectively. In order to test for a causal relationship between the mechanical properties of the pIIa-pIIb interface and Notch activation, we then studied the effect of the loss of activity of regulators of the core Notch pathway.

### Core Notch pathway components control relaxation at the daughter interface

The above results prompted us to test whether canonical Notch regulators directly influence the mechanical properties of the SOP daughter cells interface (Fig. 5A). Loss of Numb, which prevents Notch activation in pIIb cells by controlling Notch and Sanpodo trafficking (23, 24), generated a pIIa–pIIa-like interface that failed to recoil following ablation (Fig. 5B), exhibiting significantly reduced relaxation compared to control (Fig. 5D). This is associated with a membrane that becomes tortuous over time, as observed for the loss of function of AP-47 (Fig. 4C, S5A). Even though Numb is expressed in EC and SOP, these effects are restricted to SOP daughter cells (Fig. S5B). Conversely, depletion of Par3, Neuralized (Neur), or Sanpodo (Spdo), all required for Notch activation (Fig. 5A), consistently increased recoil following ablation of the SOP daughter cells interface, without affecting EC junctions mechanics (Fig. 5 C,D, S5B) and without affecting SOP shape in Spdo mutant (Fig. S5C, C’). Across multiple genetical perturbations, loss of Notch activation was associated with increased tension. These results highlighted a strong correlation between reduced Notch activity and increased tension at the interface of daughter cells, raising the question of a causal link between membrane tension and Notch signalling, which we subsequently investigated.

**Figure 5:**
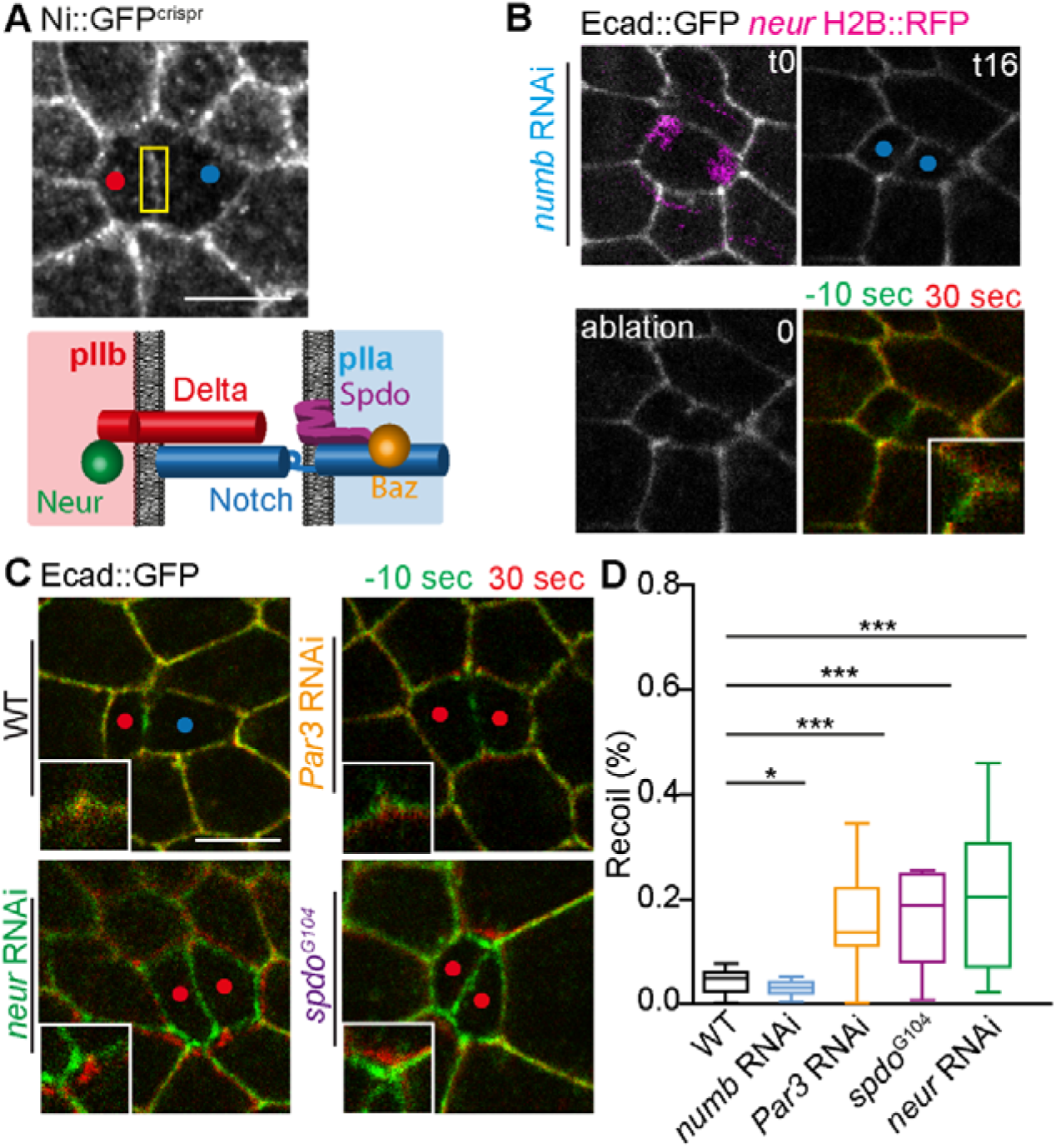
Notch cofactors regulate the mechanical properties of the SOP daughters cell interface to promote Notch differential activation. **(A)** Localization of Ni::GFP at the pIIb-pIIa interface. Ni::GFP accumulates transiently at the new interface 15 minutes after anaphase onset (adapted from (27)). (Bottom panel) Schematic summary representing the Notch signalling cascade in the pIIb/pIIa daughter cells, highlighting the distribution of Delta, Spdo, Neur, Par3, and Numb (adapted from Houssin et al., 2021). The spatial organization of Delta, Spdo, Neur, and Par3 is shown to illustrate the molecular asymmetries that accompany Notch activation. **(B)** (upper panel) Time-lapse imaging of Ecad::GFP together with neur-H2B::RFP at the pIIa–pIIa lke interface upon Numb silencing. (Lower panel) Ablation of pIIa-pIIa-like interface at 16 minutes. The t0 corresponds to the time of ablation of pIIa-pIIa-like interface. Right panel shows merges 10 seconds before ablation (green) and 30 seconds after (red). Enlarged area focused on pIIa-pIIa-like vertex. **(C)** Ablation of pIIb-pIIa (WT) and pIIb-pIIb-like interfaces upon silencing of Par3, Neuralized and Sanpodo. Green-Red images represent merges 10 seconds before ablation (green) and 30 seconds after (red). Zoomed in area focused on pIIb-pIIb-like vertices. **(D)** Quantification of Ecad recoil at the pIIb–pIIa (WT, n= 16), pIIa-pIIa-like (silencing of Numb, n= 10) and pIIb-pIIb-like (silencing of Par3, n= 8, Sanpdo, n= 11 and Neuralized,n= 18) interfaces. These results demonstrate that Notch pathway components regulate the mechanical properties of the pIIb-pIIa interface. Red and blue dots represent anterior pIIb (or pIIb-like) and posterior pIIa (or pIIa-like) cells, respectively. pIIa/pIIb cells are labelled with nuclear H2B::RFP under *neuralized* promoter. Scale bars: 5 µm. Times in minutes. *t*_₀_ = anaphase onset.

### Notch activation is sufficient to impose low interface tension

To determine the causal link between Notch activation and the properties of the membrane interface, we triggered Notch activation in SOP cells by inducing, via heat shock, the expression of the constitutively active Notch intracellular domain (Fig. 6A, Nintra)(25, 26). Expression of Nintra alone did not alter the low recoil characteristic of the pIIa–pIIb interface (Fig. 6B,B’ and Fig. 6D). However, Nintra expression in a reduced Notch activity as a result of Spdo depletion restored low recoil following ablation, reversing the increased tension induced by the loss of Spdo alone (Fig. 6B–D). This reduced restored tension is also associated with a tortuous SOP daughter cells interface characteristic of a low tense interface and a Notch gain-of-function (Fig. S6AA’) as previously observed upon depletion of Numb (Fig. S5A) and AP1 (Fig. 4C). Similar rescue by Nintra was observed upon Neur depletion (Fig. 6D). Thus, activation of Notch is sufficient to impose a low membrane tension at the SOP daughter interface. Together, our data demonstrate that the pIIa–pIIb interface is mechanically specialized and that Notch activation actively instructs its low membrane tension. Mechanical properties of the daughter interface are therefore not merely permissive for signalling but are defined by Notch activity itself.

**Figure 6:**
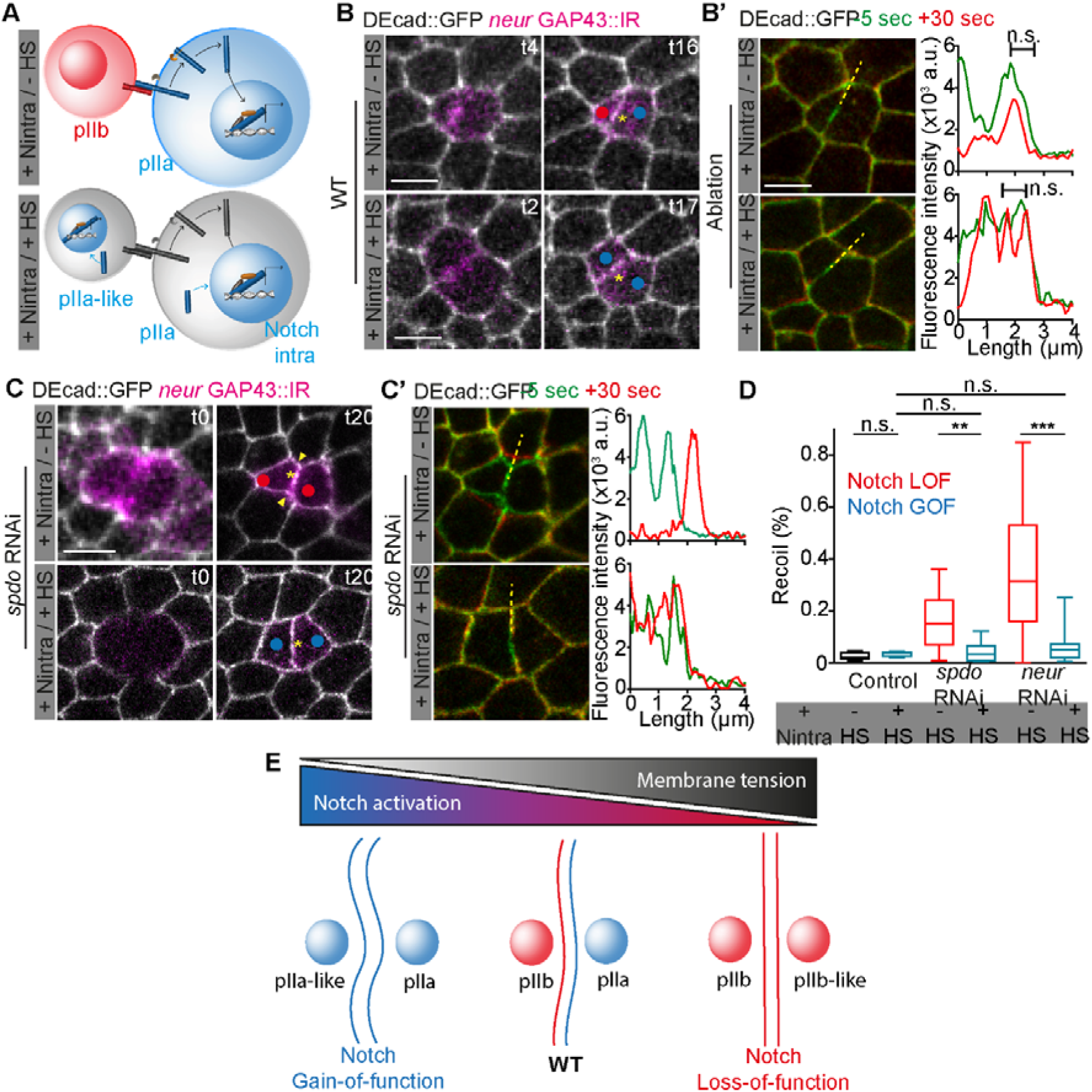
Forced Notch gain-of-function alter the mechanical properties of the pIIb-pIIa interface. (A) Schematic representation illustrating the effects of Notch gain-of-function on daughter-cell identity through Notch^intra^ expression (adapted from Pinot et al., 2024), induced by a 37°C heat-shock performed at 15h APF during 45 min. (B) Time-lapse imaging of Ecad::GFP of the pIIb-pIIa interface with and without Notch intra expression. (B’) Ablation of pIIb-pIIa (upper panel) or pIIa-pIIa-like interface (lower panel) t17 after anaphase onset. Green-Red images represent merges 5 seconds before ablation (green) and 30 seconds after (red). Fluorescence intensity profiles represent Ecad::GFP signal along the dashed white line before (green) and after ablation (red). (C) Time-lapse imaging of Ecad::GFP of the new pIIb-pIIb-like interface upon silencing of Sanpodo with and without Notch^intra^ expression. Yellow arrows represent modification of angles of pIIb-pIIb like interface characteristics of sanpodo silencing. (C’) Ablation of pIIb-pIIb-like (upper panel) or pIIa-pIIa-like interface (lower panel) t20 after anaphase onset. Green-Red images represent merges 5 seconds before ablation (green) and 30 seconds after (red). Fluorescence intensity profiles represent Ecad::GFP signal along the dashed white line before (green) and after ablation (red). (D) Recoil displacement (%) following ablation of the SOP daughter cell interface in WT (n= 10 for – HS condition, n= 8 for + HS condition), upon silencing of Sanpodo (n= 17 for – HS condition, n= 10 for + HS condition) and Neuralized (n= 10 for – HS condition, n= 19 for + HS condition) with or without Notch^intra^ expression. Blue panel represents gain-of-function phenotypes. Red panel represents loss-of-function phenotypes. Red and blue dots respectively represent anterior pIIb (or pIIb-like) and posterior pIIa (or pIIa-like) cells. SOP daughter cells are identified with GAP43::IR expressed under the *neuralized* promoter. Scale bars: 5 µm. Times in minutes. *t*_₀_ = anaphase onset. (E) A schematic diagram summarising the relationship between Notch activation and membrane tension at the apical interface of the SOP daughter cell. In Notch gain-of-function (left, blue), both cells adopt a pIIa/pIIa-like fate, correlating with a low tense and a tortuous interface. In WT (center), asymmetric Notch activation (high in pIIa, low in pIIb) correlates with a low membrane tension. In Notch loss-of-function (right, red), both cells adopt a pIIb and a pIIb-like fate, correlating with an increased tension

## Discussion

Epithelial tissues coordinate signalling and mechanics to ensure robust cell fate decisions during development. Here, we identified the pIIa–pIIb interface formed after sensory organ precursor (SOP) division as a mechanically specialized junction characterized by persistently low membrane tension (Fig. 6E). Our results demonstrate that this mechanical state is not merely permissive for signalling but is also imposed by Notch activity.

It is now recognized that mechanical traction forces are required for Notch activation, as they promote the ligand-induced, endocytosis-dependent conformational changes necessary for receptor cleavage. (Gordon, Vardar-Ulu et al. 2007; Tiyanont, Wales et al. 2011; Langridge and Struhl 2017 (6). In this framework, elevated membrane tension in the plane of AJ would negatively regulate ligand endocytosis thereby preventing ligand-induced Notch activation from the apical Notch signalling clusters. The low membrane tension measured at the AJ plane of SOP daughters is proposed here to be permissive for Notch activation. Thus, both the apical and the basal pools of Notch are present in membrane regions characterized by low membrane tension. Our findings extend this view by revealing a previously unexpected mechanism of reciprocal regulation: reducing Notch activity consistently increases junctional recoil, whereas constitutive activation of Notch restores low membrane tension even in the absence of key upstream regulators, i.e. Spdo or Neur. These results indicate that Notch signalling feeds back on junctional mechanics to define the physical properties of its own signalling interface (Fig. 6E).

The pIIa–pIIb junction displays distinctive molecular features consistent with mechanical insulation. These features include a discontinuous organisation of E-cadherin, reduced recruitment of the tension sensor Vinculin, an altered distribution of Sidekick, and a shell-like cortical arrangement of Myosin II. Together, these characteristics suggest that Notch-dependent remodelling of adhesion complexes and cortical contractility reduces effective junctional tension. We propose that such remodelling could: i) stabilize apical receptor–ligand clusters, ii) limit mechanical interference from neighbouring epidermal cells, or iii) confine signalling to a spatially defined domain.

One question does arise: why might a low-tensed membrane be advantageous for signalling? One possibility is that reduced tension stabilizes nascent signalling assemblies at the daughter interface, ensuring sustained Notch activation during the critical window of time following division. Alternatively, mechanical isolation of the interface may prevent force transmission from the surrounding epithelium, thereby buffering asymmetric fate specification against tissue-level mechanical fluctuations. In this view, Notch signalling not only determines cell identity but also establishes the mechanical context that safeguards its own activation.

More broadly, our findings support a model of coupling between signalling and mechanics, according to which developmental pathways do not merely respond to physical forces but actively instruct the mechanical identity of specific cell–cell interfaces. Such feedback may represent a general principle by which tissues coordinate morphogenesis with fate specification during asymmetric cell division.

### Limitations of the study

Our study focuses on the apical pool of Notch at adherens junctions and does not address the basal Notch pool active during cytokinesis (12), whose mechanical environment still remains poorly characterized. In our experiments, the junctional tension is inferred from laser ablation, and we use indirect readout that integrates multiple mechanical parameters. Although Notch activity correlates with reduced membrane tension, its causality remains difficult to fully disentangle, as several perturbations tested in this study affect both mechanics and signalling. In particular, we lack a perturbation that selectively increases tension without directly altering the Notch pathway. Finally, the timing of Notch-dependent nuclear responses suggests that tension changes are unlikely to arise from transcriptional outputs, raising the possibility of rapid, non-transcriptional regulation of membrane mechanics by Notch. While our findings support a bidirectional coupling between Notch signalling and mechanics in this experimental system, the extent to which this principle generalizes to other tissues, organisms, or modes of Notch activation remains to be determined.

## Materials and Methods

### Dissection for live imaging

Pupae were dissected between 16 and 17 hours APF. Placed on glass slide with double-sided tape, the pupal case was removed from the head and the dorsal thorax with microdissecting tweezers and scissors. Pillars made of 4 and 5 glass coverslips (18X18mm) were put at the anterior and posterior side of the pupae respectively. Then, a glass coverslip of 24X18mm with a thin layer of Voltalef 10S oil is placed on the pillars so that a meniscus is formed between the dorsal thorax and the coverslip (10) Cliquez ou appuyez ici pour entrer du texte.

### Microscopy

To image living pupae expressing RNAi, we used a confocal microscope ZEISS LSM980 Airyscan, equipped with 3 excitation laser diodes 488 nm, 561 nm and 635 nm. Acquisitions were made with a 63X PlanApo lens with high numerical aperture (N.A. 1.4). SOP and pIIa-pIIb daughter cells were identified and timed with the nuclear marker histone H2B::IR, histone H2B::RFP, membrane marker GAP43 expressed under the *neuralized* promoter or Par3::mScarlet asymmetrically localized in SOP during metaphase to anaphase transition (11).

### Laser-based nano-ablation

Laser ablation experiments were performed using a pulsed IR Mai-Tai HP laser from Spectra physics set to 800 nm and a laser power of 2.9 W, mounted on a Zeiss airyScan LSM 880 or Leica SP5 inversed microscopes. The laser beam was focused through an oil-immersion lens of high numerical aperture (Plan-Apochromat ×63-1.4NA). Except for Septate junction experiments, photo-ablation was performed in the focal plane at the center of bicellular adherens junctions marked with Ecad::GFP, following an area that represent a third of the total length of the ablated junction. We used a set power of 40% combined with 2 iterations.

### Image analysis

Recoil velocities (µm/s) and Recoil (%) coming from laser-based experiments were extracted through home-made Matlab program. Recoil velocities are defined as the instantaneous velocities of relaxation following ablation. Recoil final displacement in % is defined by (_Lmax following ablation_ – L _before ablation_ / L _before ablation_) where L is the length of the junction.

### Statistical analyses

Statistical differences between two conditions were evaluated by an F test followed by a Student t test using GraphPad Prism. Statistical significances were represented as follows: not significant (ns) p-value ≥ 0.05; *p-value ≤ 0.05; **p-value ≤ 0.01; ***p-value ≤ 0.001.

### Ethics

All experiments were done according to ethics rules concerning the use of *Drosophila melanogaster*, and GMOs declared in the declaration of contained use of genetically modified organisms (GMOs) of containment class 1 n °L1-560 from the Ministère de l’Enseignement Supérieur, de la Recherche et de l’Innovation.

## Acknowledgments

We would like to thank J. Januschke, T. Lecuit, F. Schweisguth, the Bloomington Drosophila Stock Center, the Vienna Drosophila Resource Center and InDroso for providing fly lines. We thank L. Chesneau, C. Dillard, J. Kubiak and C. Roubinet for critical reading. We acknowledge the engineers of the Microscopy Rennes Imaging Center (MRic, BIOSIT, Biogenouest) for assistance. MRic is a member of the national infrastructure France-BioImaging supported by the French National Research Agency (*ANR-24-INBS-0005 FBI BIOGEN*). This project was supported in part by Fondation ARC (PJA20151203510) and the ANR (ANR-20-CE13-0015) to R.L.B.

## Author contribution

**Conceptualization**: Roland Le Borgne and Mathieu Pinot. **Methodology**: Mathieu Pinot, Roland Le Borgne. **Validation**: Mathieu Pinot, Roland Le Borgne. **Formal analysis**: Mathieu Pinot. **Visualization**: Mathieu Pinot. **Writing**: Mathieu Pinot, Roland Le Borgne. **Supervision**: Roland Le Borgne. **Project administration**: Roland Le Borgne.

## Materials and Methods

### Key Ressources Table

|  | Resources | Source | Identifier |
| --- | --- | --- | --- |
| Products | Voltalef | VWR | 24627.188 |
| Gene<br>(Drosophila melanogaster) | *neur-GAP43::IR* | Houssin et al., 2021 |  |
|  | *neur-H2B-RFP* | Gomes et al., 2009 |  |
|  | *pnr-Gal4 / TM6, Tb* | BDSC | RRID: BDSC_3039 |
|  | *Par3::mScarlet* | Gift from J Januschke | FLYB:FBgn0000163 |
|  | MyoII::GFP | Daniel et al., 2018 |  |
|  | MyoII::RFP | Daniel et al., 2018 |  |
|  | Sdk::GFP | BDSC | RRID:BDSC_60169 |
|  | Ecad::GFP-GE23,<br>MyoII::RFP/ Cyo;<br>*neur-GAP43::IRFP*,<br>*pnr-Gal4 / TM6, Tb* | This study |  |
|  | hs-Notch intra ;<br>*pnr-GAL4 / SM5*,<br>TM6C | This study |  |
|  | Vinculin::GFP /<br>TM3, Ser | Gift from T. Lecuit |  |
|  | *FRT82B, nls-RFP* | BDSC | RRID:BDSC_30555 |
|  | *hsFLP* | BDSC | RRID:BDSC_6938 |
|  | *FRT82B, spdo<sup>G104</sup>* | O'Connor-Giles and Skeath, 2003 | FLYB: FBgn0260440 |
|  | *FRT82B, Sec15<sup>2</sup>* | Gift from g<br>Kriegel |  |
|  | *FRT82B, AP47<sup>she11</sup>* | Loyer et<br>al., 2015 |  |
|  | *UAS-Spdo dsRNA*<br>(II) | VDRC | <u>VDRC 104092</u> |
|  | *UAS-Nb dsRNA*<br>(III) | BDSC | FLYB: FBgn0002973;<br>RRID:BDSC 35045 |
|  | *UAS-Neur dsRNA* | BDSC | FLYB: FBgn0002932;<br>RRID:BDSC_26023 |
| Software | Matlab R2015 |  |  |
|  | Prism | GraphPad |  |
|  | ImageJ/FIJI<br>Version1.54p | Open<br>source<br>Java<br>image<br>processing<br>program | <u>https://imagej.net/Fiji</u> |

#### Drosophila genotypes

Figure 1

- A *neur*-H2B::RFP; Ecad::GFP, MyoII::RFP
- A’ apical : *neur*-H2B::RFP; Ecad::GFP, MyoII::RFP lateral : MyoII::RFP/Y ; *neur*-H2B ::IR/+ ; ATPalpha::GFP / +
- B MyoII::GFP/Y;; *pnr*-Gal4/ *neur*-GAP43::IR
- C Ecad::GFP, MyoII::RFP; *pnr*-Gal4/ *neur*-GAP43::IR
- D *neur*-H2B::RFP; Ecad::GFP, MyoII::RFP

Figure 1. Figure Supplemental 1

- A’ *neur*-H2B::RFP; Ecad::GFP, MyoII::RFP A’ MyoII::RFP/Y ;; ATPalpha ::GFP / *neur-*H2B ::IR A’’ ; neur H2B ::IR / GAP43 ::mEOSFP
- B MyoII::GFP/Y ;; *pnr*-Gal4/*neur*-GAP43::IR
- **C** *neur*-H2B::RFP; Ecad::GFP, MyoII::RFP **C’** MyoII ::GFP/Y ;; *pnr*-Gal4/neur GAP43::IR
- **D** *neur*-H2B::RFP; Ecad ::GFP, MyoII ::RFP ATPalpha::YFP ; *neur*-H2B::IR

Figure 2

- B, C, D *neur*-H2B::RFP; Ecad::GFP, MyoII::RFP

Figure 2. Figure Supplemental 2

A, B, C *neur*-H2B::RFP; Ecad::GFP, MyoII::RFP

Figure 3

- A Par::mScarlet/Y ; Ecad::GFP ; *pnr*-Gal4/+
- B *neur*-H2B::RFP; Ecad::GFP, MyoII::RFP
- C Sdk::GFP, MyoII::RFP/Y;; *neur*-GAP43 ::IR/+
- D MyoII::GFP/Y ; *neur*-H2B::IR/+
- E *neur*-H2B::IR/+; MyoII heavy chain::YFP/+

Figure 3. Figure Supplemental 3

- A neur H2B::RFP; Ecad ::GFP, MyoII ::RFP
- B Baz::mScarlet/Y ;Ecad ::GFP ; *pnr*-Gal4/+
- C Vinc::GFP / *neur*-GAP43 ::IR
- D MyoII::GFP/Y ;; *neur*-H2B::IR/+
- E ; *neur*-H2B ::IR/+; MyoII heavy chain::YFP/+

Figure 4

- B *hs*-FLP, *neur*-H2B::RFP/Y; Ecad::GFP/+;FRT82B, nls-RFP/FRT82B, *Sec15 hs*-FLP, *neur*-H2B::RFP/Y; Ecad::GFP/+;FRT82B, nls-RFP/FRT82B, AP47she11
- C *hs*-FLP, *neur*-H2B::RFP/Y; Ecad::GFP/+;FRT82B, nls-RFP/FRT82B, *Sec15*
- D *hs*-FLP, *neur*-H2B::RFP/Y; Ecad::GFP/+;FRT82B, nls-RFP/FRT82B, *AP47^she11^*

Figure 4. Figure Supplemental 4

*hs*-FLP, *neur*-H2B::RFP/Y; Ecad::GFP/+;FRT82B, nls-RFP/FRT82B, *AP47^she11^*

Figure 5

- A Ni::GFP/Y, *neur*-H2B::RFP ; *pnr-*GAL4/+
- B *neur*-H2B::RFP/Y; Ecad ::GFP, MyoII:/RFP ; *pnr-*GAL4/ Numb RNAi
- C *neur*-H2B::RFP/Y; Ecad ::GFP, MyoII:/RFP / Par3 RNAi ; *pnr*-GAL4/+ *neur*-H2B::RFP/Y; Ecad ::GFP, MyoII ::RFP/+; *pnr*-GAL4/neur RNAi *hs*-FLP, neur H2B::RFP/Y; Ecad::GFP/+;FRT82B, nls-RFP/FRT82B, spdo G104

Figure 5. Figure Supplemental 5

- A neur H2B::RFP/Y; cad ::GFP, MyoII:/RFP ; pnr-GAL4/ Numb RNAi

Figure 6

- B ; *hs-Notch intra* / Ecad::GFP, MyoII::RFP; pnr-GAL4 , neur GAP43::IR /+
- C ; *hs-Notch intra* / Ecad::GFP, MyoII::RFP, spdo RNAi; pnr-GAL4 neur GAP43::IR / +

## Supplemental Figures

**Figure S1:**
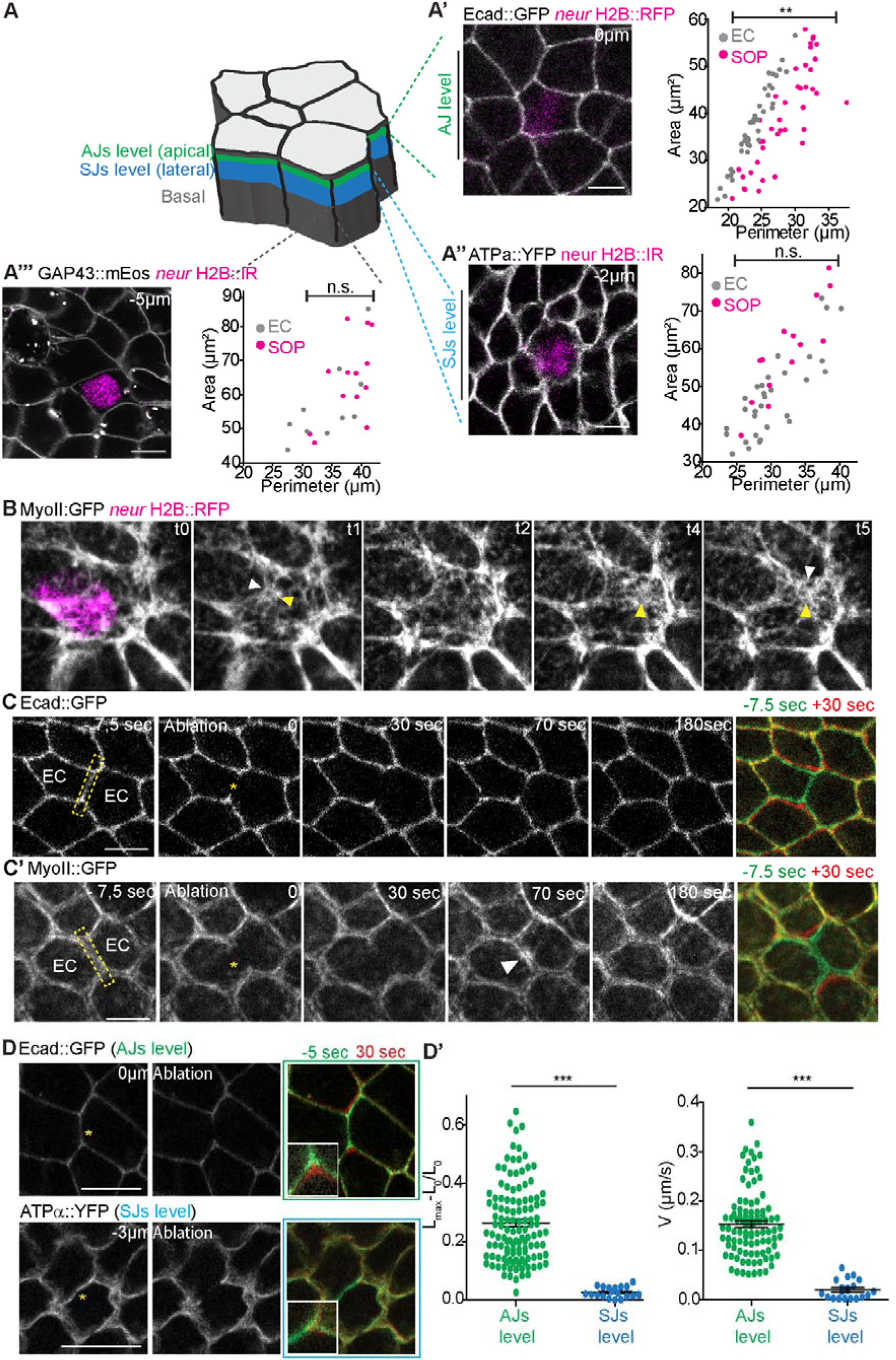
Related to Fig. 1. Interplay between atypical shape and mechanical properties of sensory organ precursors (SOPs). **(A)** Schematic representation of the monolayer epithelium of the *Drosophila* pupal notum. Adherens (AJs) and septate (SJs) junctions are represented in green and blue, respectively. **(A’)** Confocal image of SOP expressing Ecad::GFP. Plot of cell perimeter in function of cell area at the level of AJs (n = 42 for SOPs and n = 71 for ECs). **(A’’)** Confocal image of SOP expressing ATPa::YFP. Plot of cell perimeter in function of cell area at the level of SJs (n = 14 for SOPs and n = 23 for ECs). **(A’’’)** Confocal image of SOP expressing GAP43::mEos. Plot of cell perimeter in function of cell area at the lateral membrane (n = 13 for SOPs and n = 25 for ECs). In (A’-A’’-A’’’), grey dots represent ECs and black dots represent SOPs. SOPs are identified with the nuclear marker histone H2B::IR expressed under the *neuralized* promoter. **(B)** Time-lapse imaging of MyoII::GFP in SOP marked with the nuclear marker H2B::RFP (magenta). White arrows represent the curved interface. Yellow arrows represent the local accumulation of medial MyoII closed to the interface. (C-C’) Time-lapse imaging following the ablation of EC-EC interface marked with Ecad::GFP (C) or MyoII::GFP (C’). Right panel represents the merge of two pictures, one taken 7.5 seconds before ablation (green) and the other one taken 30 seconds after ablation. **(D)** Confocal images representing ablation of EC/EC junction at the level of AJs (upper panel) and SJs (lower panel). Left panels represent pictures cells before ablation. Middle panels represent ablation time point. Right panel represent the merge of two pictures, one taken 5 seconds before ablation (green) and the other one taken 30 seconds after ablation. White squares represent the magnification performed on a vertex. The difference between green and red channels are related to the junction tension. **(D’)** Quantification of the recoil maximal displacement (%) and the recoil velocity (µm/s) following ablation of the EC-EC junction at the levels of AJs (green dots n = 121) and SJs (blue dots n = 21). Scale bars are 5 µm. Times are expressed in seconds in (C, D) and in minutes in (B). Yellow stars represent ablated junction.

**Figure S2:**
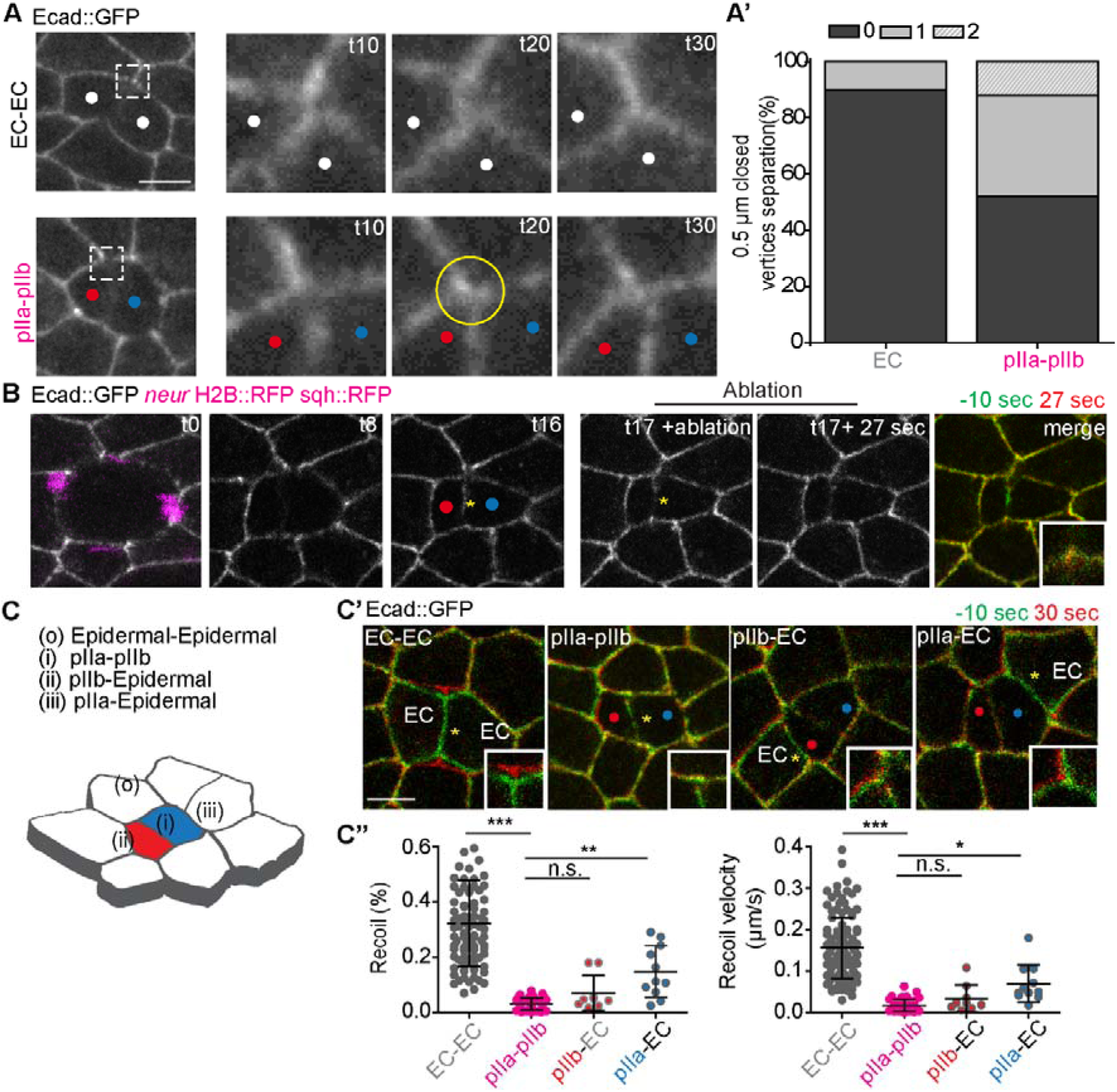
Related to Fig. 2: pIIa-pIIb adhesive interface exhibits low membrane tension. **(A)** Evolution of the formation of the new vertices during EC/EC cytokinesis (upper panel) or during SOP cytokinesis (lower panel). (right panels) Magnification of the formation of the new vertex presented on panel (A). The formation of the vertices leads to a symmetrical arrangement during EC cytokinesis and to a configuration in which the vertices are very close together during SOP cytokinesis. (A’) Quantification of the distribution of the close-vertices configuration during EC (n = 26) and SOP (n = 38) cytokinesis. A closely spaced vertices configuration is defined when two adjacent vertices are spaced less than 0.5 µm from each other. **(B)** Experimental procedure to perform ablation at the pIIa/pIIb interface at a specific time points after anaphase onset. First a time-lapse imaging is performed with Ecad::GFP marker. Between t15 and t20 after the onset of anaphase, ablation is performed at the pIIa-pIIb interface. The panel on the far right shows a composite of two images taken during the ablation process: one taken 10 seconds before the ablation (in green) and the other taken 27 seconds after the ablation (in red). White squares represent the magnification performed on a vertex before and after ablation. A separation of green and red channels means that the junction followed a relaxation process after ablation whereas a colocalization of green and red channels means the absence of a relaxation process following ablation characteristic of a low-tense interface. **(C)** 3D-schematic representation of *Drosophila* pupal notum composed of ECs (white cells) and pIIb (red) / pIIa (blue) cells. (C’) Ablation of EC-EC, pIIb-pIIa, pIIb-EC, pIIa-EC interfaces, respectively. Images represent the merge of two pictures taken 10 seconds before ablation (green) and 30 seconds after ablation (red). (C’’) Quantification of the recoil maximal displacement (%) and recoil velocity (µm/s) following ablation of EC-EC (n = 148), pIIa-pIIb, (n = 47), pIIb-EC (n = 16) and pIIa-EC (n = 11) interfaces. Scales bars are 5 µm. Red and blue dots represent anterior pIIb and posterior pIIa cells, respectively. Times are expressed in minutes.

**Figure S3:**
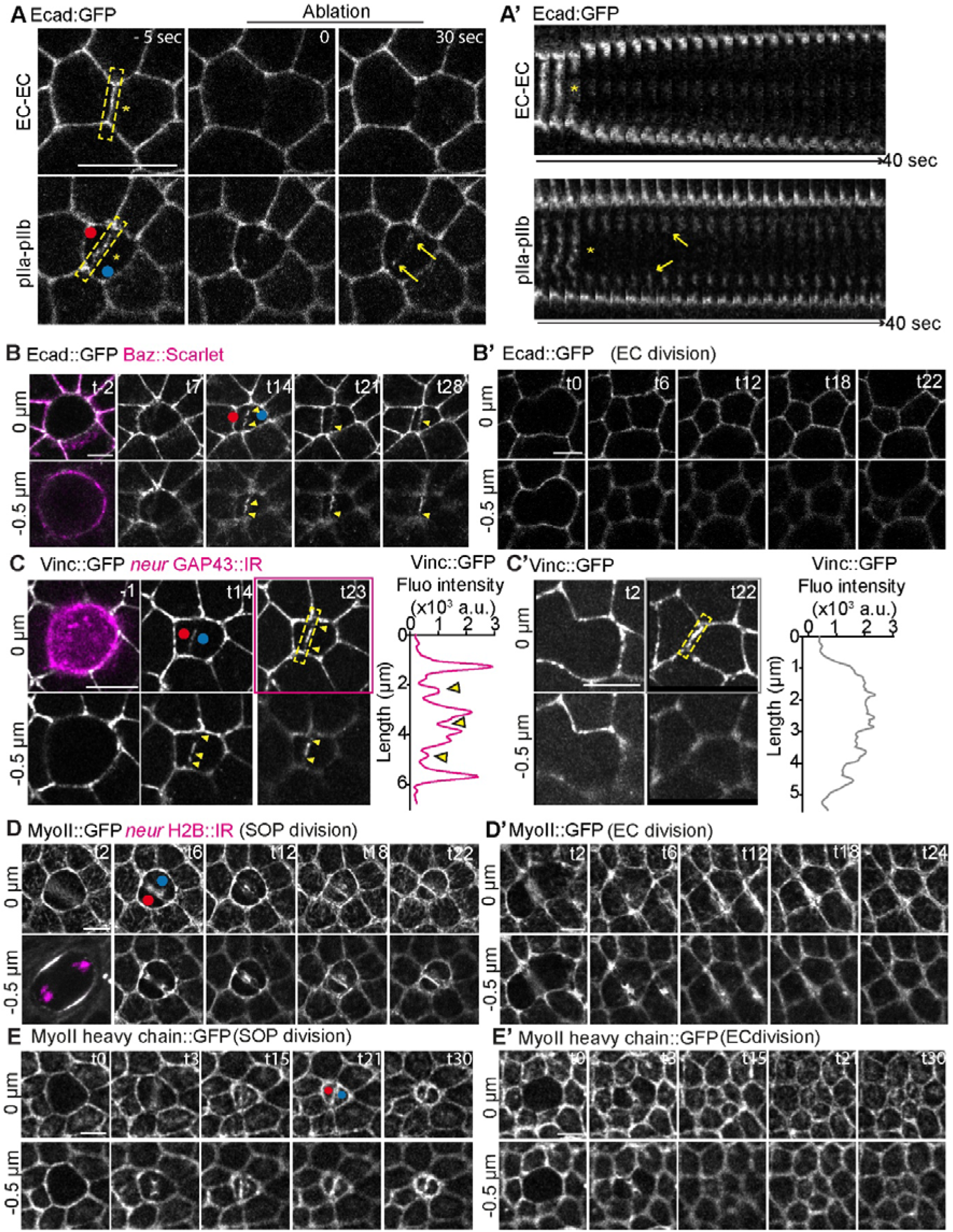
Related to Fig. 3: Distribution and dynamics of adhesive markers during SOP division. **(A)** Two examples of ablation of EC-EC junction and the pIIa-pIIb interface. The left panels represent cells before ablation. The middle panels illustrate the ablation of the junction. The right panels show cells 30 seconds after ablation. (A’) Pseudo kymographs are generated from the dashed yellow rectangles in (A). Kymographs represent the time-evolution of the ablated EC-EC or pIIa-pIIb junction. Yellow stars represent ablated junction. Yellow arrows point to the remaining amount of Ecad::GFP at the ablated junction. We noticed that this remaining amount is stabilized over time following ablation of the pIIa-pIIb junction. **(B-B’)** Time-lapse imaging of Ecad::GFP during SOP (n= 20) and EC cytokinesis (n= 20) at AJ level and 0.5 µm below. SOP daughter cells identified by Par3::mScarlet. Related to figure 3. **(C-C’)** Time-lapse imaging of Vinculin::GFP during SOP (n= 4) and EC cytokinesis at the level and 0.5 µm below AJs. Plot profiles represent the yellow dashed area shown in (C). Yellow arrowheads represent Vinc::GFP discontinuities. **(D-D’)** Time-lapse imaging of MyoII::GFP during SOP (n= 10) and EC cytokinesis at AJ level and 0.5 µm below. Related to Figure 3. **(E-E’)** Time-lapse imaging of MyoII heavy chain::GFP during SOP (n= 10) and EC (n= 8) cytokinesis at AJ level and 0.5 µm below. Related to Figure 3. Red and blue dots correspond to anterior pIIb and posterior pIIa cells, respectively. pIIa/pIIb cells marked by nuclear H2B::RFP under *neuralized* promoter. Scale bars: 5 µm. Times are in minutes.

**Figure S4:**
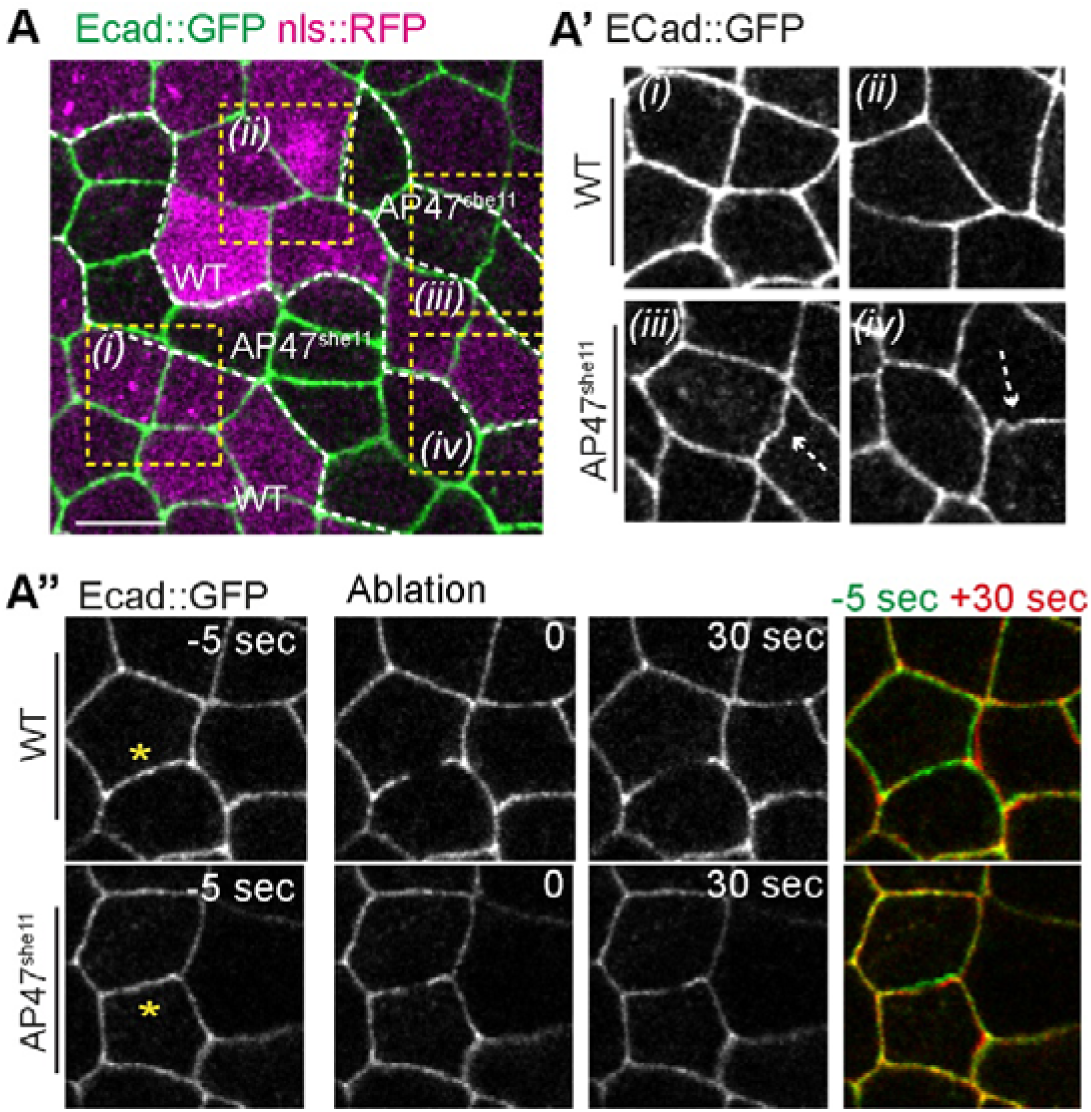
related to Fig. 4: Vesicular trafficking regulators control mechanical properties of the newly formed EC-EC interface. (A-A’) Confocal image of Ecad::GFP expressed in *AP47^she11^* mutant clones and WT cells. (A’) Tortuous interfaces observed in epidermal cells in AP47*^she11^* conditions (iii) and (iv) and WT (i) and (ii) conditions. (A’’) Ablation of epidermal cells in WT conditions and upon silencing of AP47. Time-lapse imaging of ablations. Right images represent merge of two pictures, one taken 5 seconds before ablation (green) and the second 30 seconds after ablation (red). Scale bars, 5 µm.

**Figure S5:**
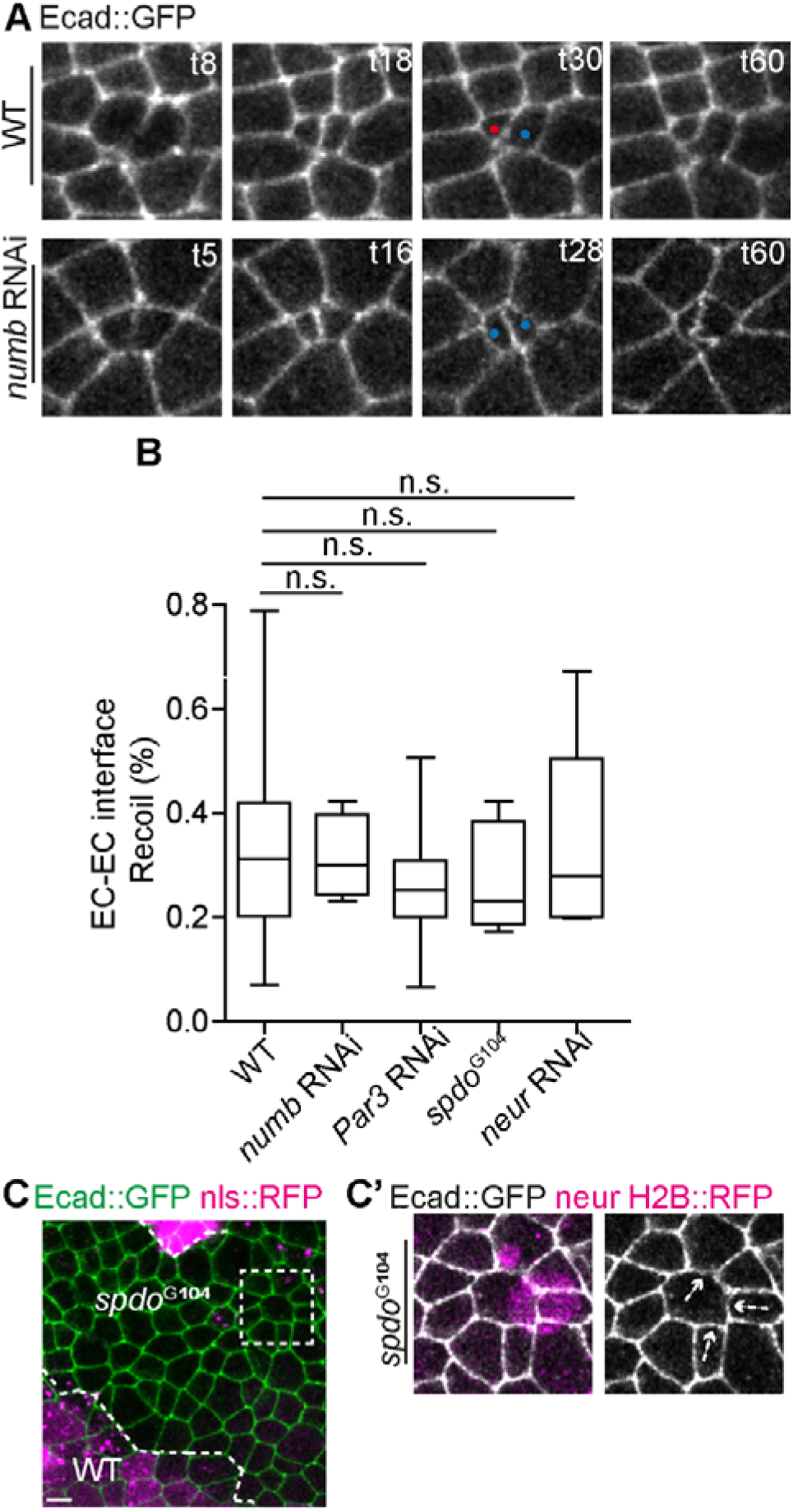
related to Fig. 5: Notch cofactors regulate the mechanical properties of the SOP daughters cell interface to promote Notch differential activation. (A) Time-lapse imaging of Ecad::GFP in WT (upper panel) and upon silencing of Numb (lower panel). Upon loss of Numb, pIIa-pIIa-like interface became highly tortuous compared to WT pIIa-pIIb interface. (B) Recoil maximal displacement (%) following ablation of EC-EC interfaces in WT conditions (n= 148) or upon silencing of numb (n= 8), Par3 (n= 22), sanpodo (n= 5) and neuralized (n= 6). Red and blue dots correspond to anterior pIIb and posterior pIIa (or pIIa-like) cells, respectively. Scale bars, 5 µm. Time is expressed in minutes. (C-C’) SOP shape in *spdo^G104^* conditions. Dashed white arrows point to curved interface in SOP-EC interface as presented in WT conditions (Figure 1)

**Figure S6:**
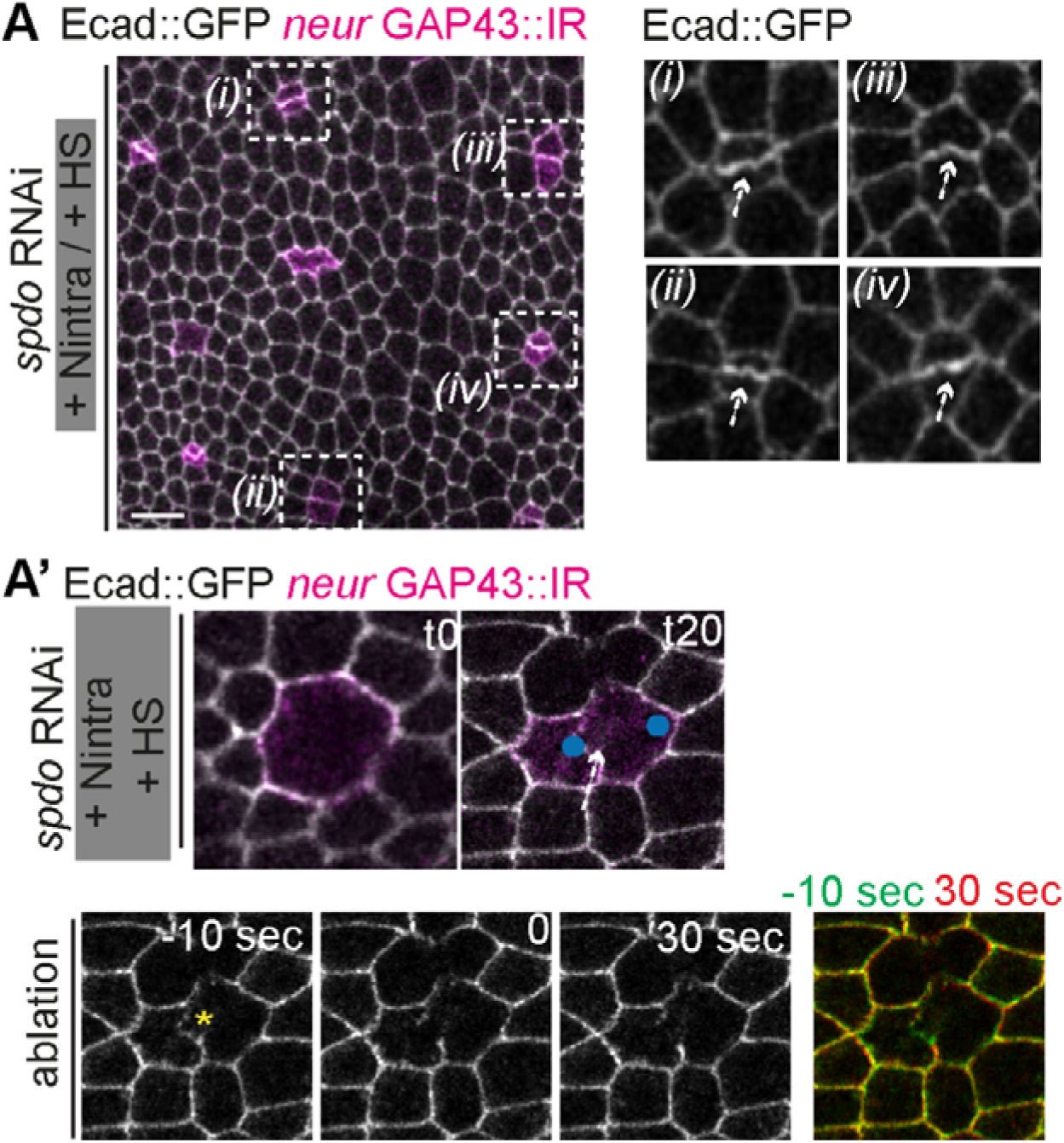
related to Fig. 6: Forced Notch gain-of-function alter the mechanical properties of the pIIb-pIIa interface. (A) Time-lapse imaging of Ecad::GFP in Notch intra condition upon silencing of *spdo*. pIIa-pIIa-like interface are tortuous compared to WT pIIa-pIIb interface. (A’) Time-lapse imaging of Ecad::GFP of the pIIa-pIIa interface upon silencing of spdo and activation of Notch intra revealed a tortuous interface 20 minutes after anaphase onset. Ablation of the pIIa-pIIa interface (related to figure 6). Green-Red images represent merges 10 seconds before ablation (green) and 30 seconds after (red).

## References

1. Kumar A, Placone JK, & Engler AJ (2017) Understanding the extracellular forces that determine cell fate and maintenance. Development 144(23):4261–4270.

2. Alvarez Y & Smutny M (2022) Emerging Role of Mechanical Forces in Cell Fate Acquisition. Frontiers in Cell and Developmental Biology Volume 10–2022.

3. Gordon WR, et al. (2007) Structural basis for autoinhibition of Notch. (1545-9993 (Print)).

4. Tiyanont K, et al. (2011) Evidence for increased exposure of the Notch1 metalloprotease cleavage site upon conversion to an activated conformation. (1878-4186 (Electronic)).

5. Langridge PD & Struhl G (2017) Epsin-Dependent Ligand Endocytosis Activates Notch by Force. (1097-4172 (Electronic)).

6. Kasirer S & Sprinzak D (2024) Interplay between Notch signaling and mechanical forces during developmental patterning processes. Current opinion in cell biology 91:102444.

7. Priya RA-O, et al. (2020) Tension heterogeneity directs form and fate to pattern the myocardial wall. (1476-4687 (Electronic)).

8. Blackie L, et al. (2021) A combination of Notch signaling, preferential adhesion and endocytosis induces a slow mode of cell intercalation in the Drosophila retina. LID - 10.1242/dev.197301 [doi] LID - dev197301. (1477-9129 (Electronic)).

9. Schweisguth F (2015) Asymmetric cell division in the Drosophila bristle lineage: from the polarization of sensory organ precursor cells to Notch-mediated binary fate decision. WIREs Developmental Biology 4(3):299–309.

10. Gho M, Bellaïche Y Fau - Schweisguth F, & Schweisguth F (1999) Revisiting the Drosophila microchaete lineage: a novel intrinsically asymmetric cell division generates a glial cell. (0950-1991 (Print)).

11. Houssin E, Pinot MA-O, Bellec KA-O, & Le Borgne RA-OX (2021) Par3 cooperates with Sanpodo for the assembly of Notch clusters following asymmetric division of Drosophila sensory organ precursor cells. LID - 10.7554/eLife.66659 [doi] LID - e66659. (2050-084X (Electronic)).

12. Trylinski M, Mazouni K, & Schweisguth F (2017) Intra-lineage Fate Decisions Involve Activation of Notch Receptors Basal to the Midbody in <em>Drosophila</em> Sensory Organ Precursor Cells. Current Biology 27(15):2239–2247.e2233.

13. Bellec K, Gicquel I, & Le Borgne R (2018) Stratum recruits Rab8 at Golgi exit sites to regulate the basolateral sorting of Notch and Sanpodo. Development 145(13).

14. Durel E, et al. (2025) Regulation of cell shape and mechanics by Rho GEFs and GAPs in a proliferative epithelial tissue. LID - jcs264163 [pii] LID - 10.1242/jcs.264163 [doi]. (1477-9137 (Electronic)).

15. Heisenberg CP & Bellaïche Y (2013) Forces in tissue morphogenesis and patterning. (1097-4172 (Electronic)).

16. Couturier L, et al. (2017) Regulation of cortical stability by RhoGEF3 in mitotic Sensory Organ Precursor cells in Drosophila. (2046-6390 (Print)).

17. Higashi T, Arnold Torey R, Stephenson Rachel E, Dinshaw Kayla M, & Miller Ann L (2016) Maintenance of the Epithelial Barrier and Remodeling of Cell-Cell Junctions during Cytokinesis. Current Biology 26(14):1829–1842.

18. Langevin J, et al. (2005) Drosophila exocyst components Sec5, Sec6, and Sec15 regulate DE-Cadherin trafficking from recycling endosomes to the plasma membrane. (1534-5807 (Print)).

19. Jafar-Nejad H, et al. (2005) Sec15, a component of the exocyst, promotes notch signaling during the asymmetric division of Drosophila sensory organ precursors. (1534-5807 (Print)).

20. Benhra N, et al. (2011) AP-1 controls the trafficking of Notch and Sanpodo toward E-cadherin junctions in sensory organ precursors. (1879-0445 (Electronic)).

21. Loyer N, Kolotuev I, Pinot M, & Le Borgne R (2015) Drosophila E-cadherin is required for the maintenance of ring canals anchoring to mechanically withstand tissue growth. (1091-6490 (Electronic)).

22. Bellec K, Pinot M, Gicquel I, & Le Borgne RA-OX (2021) The Clathrin adaptor AP-1 and Stratum act in parallel pathways to control Notch activation in Drosophila sensory organ precursors cells. LID - 10.1242/dev.191437 [doi] LID - dev191437. (1477-9129 (Electronic)).

23. Cotton M, Benhra N Fau - Le Borgne R, & Le Borgne R (2013) Numb inhibits the recycling of Sanpodo in Drosophila sensory organ precursor. (1879-0445 (Electronic)).

24. Couturier L Fau - Mazouni K, Mazouni K Fau - Schweisguth F, & Schweisguth F (2013) Inhibition of Notch recycling by Numb: relevance and mechanism(s). (1551-4005 (Electronic)).

25. Remaud S, Audibert A, & Gho M (2008) S-Phase Favours Notch Cell Responsiveness in the Drosophila Bristle Lineage. PLOS ONE 3(11):e3646.

26. Le Borgne R, Bellaïche Y Fau - Schweisguth F, & Schweisguth F (2002) Drosophila E-cadherin regulates the orientation of asymmetric cell division in the sensory organ lineage. (0960-9822 (Print)).

27. Pinot MLB, R. (2024) Spatio-Temporal Regulation of Notch Activation in Asymmetrically Dividing Sensory Organ Precursor Cells in Drosophila melanogaster Epithelium. LID - 10.3390/cells13131133 [doi] LID - 1133. (2073-4409 (Electronic)).

